# *Ex Vivo* Biomechanical Characterization of Umbilical Vessels: Possible Shunts in Congenital Heart Palliation

**DOI:** 10.1101/2023.01.07.523118

**Authors:** S-I. Murtada, A.B. Ramachandra, J.D Humphrey

## Abstract

Children born with single ventricle defects undergo staged palliative surgeries to reconstruct the circulation to improve transport of deoxygenated blood to the lungs. As part of the first surgery, a temporary shunt (Blalock-Thomas-Taussig) is often created in neonates to connect a systemic and a pulmonary artery. Standard-of-care shunts are synthetic, which can lead to thrombosis, and much stiffer than the two host vessels, which can cause adverse mechanobiological responses. Moreover, the neonatal vasculature can undergo significant changes in size and structure over a short period, thus constraining the use of a non-growing synthetic shunt. Recent studies suggest that autologous umbilical vessels could serve as improved shunts, but there has not been a detailed biomechanical characterization of the four primary vessels – subclavian artery, pulmonary artery, umbilical vein, and umbilical artery. Herein, we biomechanically phenotype umbilical veins and arteries from prenatal mice (E18.5) and compare them to subclavian and pulmonary arteries harvested at two critical postnatal developmental ages (P10, P21). Comparisons include age-specific physiological conditions and simulated ‘surgical-like’ shunt conditions. Results suggest that the intact umbilical vein is a better choice as a shunt than the umbilical artery due to concerns with lumen closure and constriction related-intramural damage in the latter. Yet, decellularization of umbilical arteries may be a viable alternative, with the possibility of host cellular infiltration and subsequent remodeling. Given recent efforts using autologous umbilical vessels as Blalock-Thomas-Taussig shunts in a clinical trial, our findings highlight aspects of the associated biomechanics that deserve further investigation.

## Introduction

Infants born with a single ventricle defect typically undergo three staged surgeries (Norwood, Glenn, Fontan) over a 2-to-3-year period to reconstruct the circulatory system, culminating in a circuit in which deoxygenated blood from the vena cava flows directly to the pulmonary arteries without cardiac assistance. These stages of reconstruction depend on the type of congenital heart defect, but the earliest can include a Blalock-Thomas-Taussig (BT) shunt wherein a graft is typically placed between a subclavian (systemic) and pulmonary artery. Synthetic grafts, often PTFE (polytetrafluoroethylene), are used despite associated risks including thrombosis, which occurs in up to 20% in these newborns (Hoganson et al., 2018). There is, therefore, a need for better alternatives.

Umbilical vessels represent promising vascular conduits, especially in pediatric patients wherein growth is critical (Vlessis et al., 1995). A recent clinical trial (NCT02766998) explored the possibility of using the autologous umbilical vein rather than a synthetic graft as a BT shunt. Preliminary results are promising (https://discoveries.childrenshospital.org/having-faith-a-novel-approach-to-heart-surgery/), including reduced thrombosis and adequate hemodynamics until the Glenn procedure, typically 4-6 months later. Nevertheless, important metrics remain unknown; there is limited information on the relative geometric, material, and structural properties of the different vascular regions involved in a BT shunt procedure during early postnatal development. Using a murine model, we recently identified key developmental times, from perinatal to late postnatal, at which the aorta undergoes critical changes in hemodynamic loading, structure, and function (Murtada et al. 2021). Herein, we used the same methods to quantify biomechanical properties and function of late embryonic (E18.5) umbilical veins (UV) and umbilical arteries (UA) and compared them to those of right subclavian arteries (RSA) and pulmonary arteries (RPA) from healthy wild-type mice at two critical development times (postnatal days P10 and P21), during which large arteries undergo significant changes. We also computed possible acute mechanical changes in these four vessels for a simulated creation of an early (P10) or later (P21) BT shunt (‘surgical-like’ condition). Finally, we compared basic geometric and structural changes in UV and UA following decellularization, given their potential for use as autologous grafts (Gui et al., 2009). These data provide further insight into the possible translatability of autologous umbilical vessels as shunts in congenital heart surgeries.

## Methods

### Vessel Isolation

All live mouse procedures were approved by the Institutional Animal Care and Use Committee at Yale University. Following euthanasia, the umbilical cord was separated from the placenta and excised from C57BL/6J mice at gestational day E18.5. Umbilical vessels were delineated by opening the abdomen of each fetus and locating the UV via its connection to the inferior vena cava through the ductus venosus and an UA via its connection to an iliac artery; both were excised by precision micro-dissection. The lumen of the intact UA closed immediately after separating the umbilical cord from the placenta and fetus, as would occur naturally at birth (Nandadasa et al., 2020). Separate groups were used for biomechanically phenotyping native and decellularized umbilical vessels. Following euthanasia of additional non-pregnant C57BL/6J mice, RSA and RPA were excised at either P10 or P21 and prepared for ex vivo testing by gently removing excessive perivascular tissue and ligating branches, if any. All intact cylindrical vessels (UV, UA, RSA, RPA) were cannulated on custom-drawn glass micropipettes, secured with silk sutures, and mounted within a custom computer-controlled biaxial device for biomechanical testing (Ferruzzi et al., 2013) within hours of euthanasia.

### Decellularization

The 4-day decellularization process was adapted from a human umbilical artery protocol (Han et al., 2021). Briefly, on the day of harvest (day 1) isolated umbilical vessels were submerged in 8 mM CHAPS buffer solution and placed on a rocking shaker overnight at room temperature. On day 2, vessels were rinsed 3x with a phosphate buffered saline (PBS) solution and placed on a rocking shaker for 15 min; this was repeated 3x over 1 hour. The vessels were then transferred to 1.8 mM SDS buffer solution and placed in a 37^°^C water bath overnight. On day 3, vessels were rinsed 3x in PBS and placed on a rocking shaker for 1 hour; this process was repeated 8x followed by placement on the rocking shaker overnight at room temperature. On day 4, the rinsing process was repeated and the vessels were stored in 4°C PBS until testing the following day.

### Biomechanical Testing

Vessels subjected to biaxial testing (*n*=4-6 per group) were submerged in a bicarbonate-buffered Krebs-Ringer solution (Krebs) that was oxygenated with 95% O_2_ and 5% CO_2_ while kept at 37^°^C, as is standard in vascular testing (Humphrey, 2013). This procedure prevented extended lumen closure in UA, which were washed up to 3x with fresh Krebs until the lumen distended fully. UV also constricted after excision, but to a lesser degree; they dilated spontaneously after mounting within the testing device. All vessels were first subjected to isobaric and axially isometric vasoconstrictions by adding 100 mM KCl to the bathing solution at vessel-specific pressures: E18.5 UA and UV at 5 and 25 mmHg, P10 RPA at 5 mmHg, P10 RSA at 30 mmHg, P21 RPA at 7.5 mmHg, and P21 RSA at 50 mmHg (Table 1). The gross morphology and transmural structure of each umbilical vessel was monitored during stages of vasoconstriction using an optical coherence tomography (OCT) system (Callisto Model, Thorlabs, Newton, NJ). Umbilical vessels that were damaged during cannulation or compromised after two contraction tests were not used for subsequent testing.

**Table 1.** Pressures, in mmHg, imposed during isobaric-axially isometric experimental measurement of vasoactive responses (Active Test) as well as those used to compute representative passive mechanical properties for native versus two simulated surgical conditions. The passive properties were computed based on experimental measurements conducted during cyclic testing over a range of pressures and axial forces (see Methods). The subclavian and pulmonary arteries were excised from the right side and tested at two different postnatal ages: P10 and P21. The umbilical vessels were excised and tested at embryonic day E18.5. All biomechanical testing was completed within hours of euthanasia and vessel harvest; there was no attempt to preserve the umbilical vessels for the 10 to 21 days that would be needed clinically, hence all results represent the ideal case wherein umbilical properties would have been preserved.

| **Vessel** | **Active Test**<br>**(mmHg)** | **Native Condition**<br>**(mmHg)** | **Early-Surgical: P10**<br>**(mmHg)** | **Late-Surgical: P21**<br>**(mmHg)** |
| --- | --- | --- | --- | --- |
| **Subclavian Artery** | 30 (P10); 50 (P21) | 56 (P10); 90 (P21) | 56 | 90 |
| **Pulmonary Artery** | 5 (P10); 7.5 (P21) | 11 (P10); 17 (P21) | 15 | 20 |
| **Umbilical Artery** | 5 and 25 | 30 | 15 | 20 |
| **Umbilical Vein** | 5 and 25 | 5 | 15 | 20 |

Next, vessels were submerged in a Ca^++^-free Krebs solution and washed 3x to ensure passive behaviors. With axial stretch held fixed at the vessel-specific *in vivo* value, that is, the stretch at which axial force is decoupled from pressure (van Loon et al., 1977; Weizsäcker et al., 1983), the vessels were preconditioned via four cycles of pressurization: UA from 0-40 mmHg, UV from 0-20 mmHg, P10 RPA from 0-25 mmHg, P10 RSA from 0-60 mmHg, P21 RPA from 0-30 mmHg, and P21 RSA from 0-90 mmHg. Next, the vessels were subjected to a series of seven biaxial protocols: cyclic pressurization at three values of axial stretch (95%, 100%, 105% of in vivo value), and cyclic axial stretching at four fixed values of luminal pressure (e.g., 2, 5, 15, and 25 mmHg for P10 RPA and 1.5, 7.5, 15, and 20 mmHg for P21 RPA). These combined active, then passive biaxial biomechanical assessments were completed within hours of euthanasia.

### Passive Material Properties

The biaxial data were used to estimate, via nonlinear regression, best-fit values of eight material parameters within a validated four-fiber family constitutive relation (Ferruzzi et al., 2013). This relation allows one to quantify elastic energy storage *W*(defined per unit volume) and thereby to compute via standard formulae the circumferential and axial wall stress and material stiffness at any biaxial deformation (*λ*_*θ*_,*λ*_*z*_). This energy is

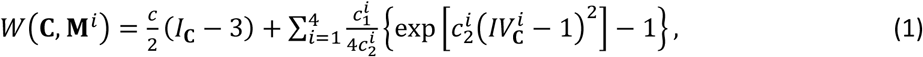

where *c* (kPa),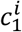, and 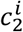 are model parameters (*i* = 1,2,3,4 denote axial, circumferential, and two symmetric-diagonal fiber family directions, respectively), with 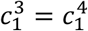 and 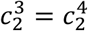 by definition. *I*_**C**_ = *tr*(**C**) and 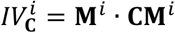 are coordinate invariant measures of the finite deformation, with **C** = **F**^**T**^**F** the right Cauchy-Green tensor computed from the deformation gradient tensor **F** = diag[*λ*_*r*_,*λ*_*θ*_,*λ*_*z*_],with *det***F** = 1 assuming incompressibility and *λ*_*r*_, *λ*_*θ*_,*λ*_*z*_ denoting principal stretches in radial, circumferential, and axial directions. 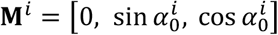 captures fiber-family directions, with 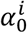 defined relative to the axial direction in the traction-free reference configuration: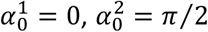, and 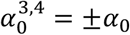 (the eighth model parameter). Physical components of the Cauchy stress tensor were computed from *W*using standard relations (Humphrey, 2013). Mean (radially averaged) values of the in-plane components of the Cauchy stress were also determined from measurable quantities,

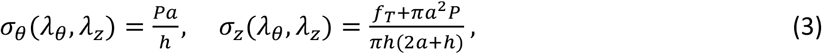

where *P* and *f*_*T*_ are transmural pressure and applied axial force, each measured with standard transducers, *a* the deformed inner radius, and *h* the deformed thickness. The radial component of stress is much less than the in-plane components, and neglected. Both *a* and *h* were calculated from incompressibility using the unloaded volume coupled with online measurements of outer diameter and axial length (Ferruzzi et al., 2013), which allowed calculation of biaxial stretches (*λ*_*θ*_,*λ*_*z*_) using standard formulae. Components of the material stiffness tensor, linearized about a configuration defined by a fixed *in vivo* pressure and axial stretch, were computed as before (Baek et al., 2007). Given the constitutive description determined over a range of pressures and axial extensions, values of key passive mechanical metrics were computed at native and simulated surgical conditions (Table 1).

### Histology

Following biomechanical testing, vessels were fixed in 10% neutral buffered formalin for 24 hours, then stored in ethanol. Subsequently, vessels were embedded in paraffin and cut into 5-μm thick transverse sections that were mounted on a glass slide and stained with Hematoxylin and Eosin (H&E), Verhoeff-Van Gieson (VVG), or Movat pentachrome (MOV). Sections were imaged using an Olympus BX/51 microscope equipped with a DP70 digital camera.

### Statistics

An unpaired non-parametric t-test (Mann-Whitney test) was used to compare results across native and surgery-like conditions, with *p* < 0.05 considered significant. All data are presented as mean ± standard error of the mean (SEM).

## Results

Figure 1 shows images of four representative specimens – RSA and RPA at P10, UV and UA at E18.5 – during biaxial testing, along with mean values of outer radius and wall thickness at representative vessel-specific values of distending pressure and axial stretch. Additional metrics for E18.5 UV and UA were estimated at representative physiological pressures (5 and 30 mmHg, respectively) and compared to those for RSA and RPA at postnatal days P10 (56 mmHg and 11 mmHg, respectively) and P21 (90 mmHg and 17 mmHg, respectively) based on reported region- and age-specific pressures (Ramachandra and Humphrey, 2019; Murtada et al. 2020; Nandadasa et al., 2020; Table S1), noting that many of these metrics were computed using best-fit constitutive parameters (Table S2). Overall structural (i.e., pressure-inner radius) behaviors exhibited by E18.5 UV and UA differed from those of P10 RSA and RPA (Fig 2A), with RSA and RPA more distensible though with the pressurized luminal radius of UV similar to those of RPA. Yet, the overall material (circumferential stress-stretch) behavior of UV and UA ranged between those of P10 RSA and RPA (Fig 2B).

**Fig 1.**
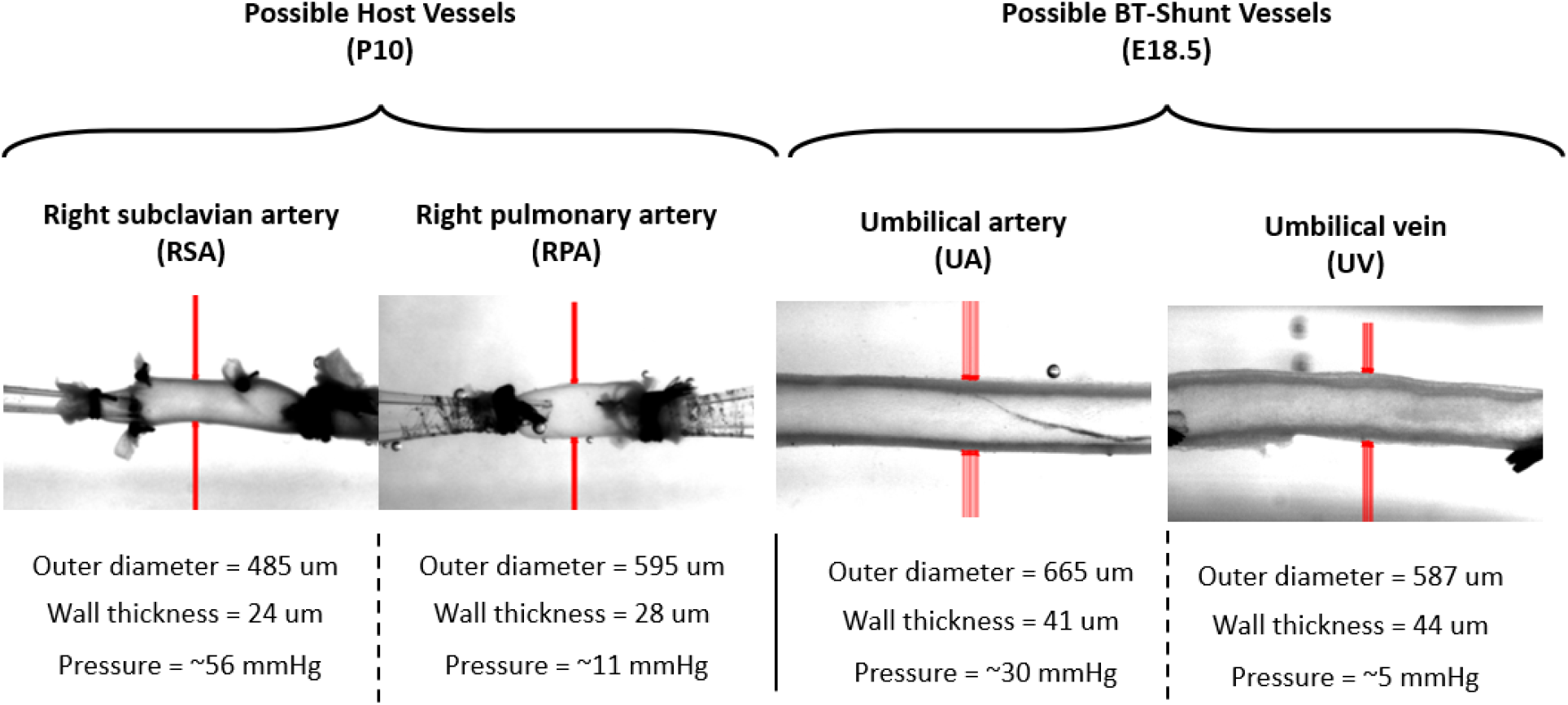
Visual comparisons of the four vessels of interest under physiologically relevant ex vivo testing conditions: right subclavian artery (RSA) and right pulmonary artery (RPA), both excised at postnatal day P10, and umbilical artery (UA) and umbilical vein (UV), both excised at embryonic day E18.5. The red vertical lines show automatic edge detection of outer diameter when held at the specimen-specific in vivo axial stretch and cyclically pressurized. Also provided are values of outer diameter and wall thickness at the in vivo relevant stretch and representative pressure.

**Fig 2.**
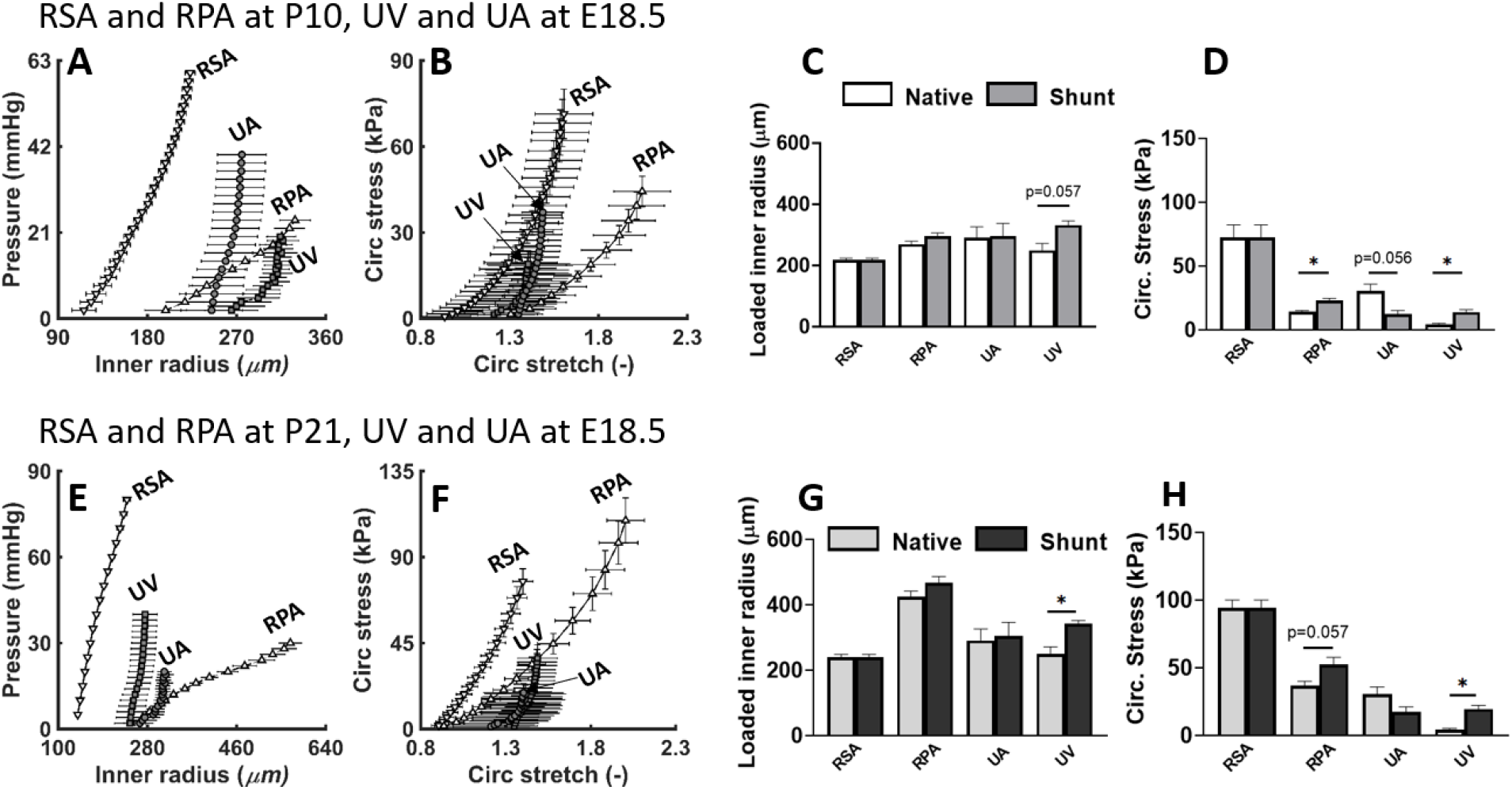
Comparison of regional passive biomechanical behaviors of the right subclavian artery (RSA, ▽) and right pulmonary artery (RPA, Δ) as well as the umbilical vein (UV, □) and umbilical artery (UA, ○). **A, B:** Pressure-inner radius and circumferential stress-stretch relationships at P10 for the RSA and RPA, with relationships for the UV and UA at E18.5 shown for comparison. **E, F:** Similar to panels A, B except at P21 for the RSA and RPA, with values for the UV and UA at E18.5 again shown for comparison. **C, D**: Values of inner radius and circumferential Cauchy stress for vessel-specific physiological loading conditions for the RSA and RPA at P10 and the UV and UA at E18.5 (white bars) as well as at simulated (surgical-like) loading conditions for an early stage (P10) BT shunt procedure (dark grey). **G, H**: Similar to panels C, D except for the RSA and RPA at P21 and the UV and UA E18.5 (light grey) as well as at simulated (surgical-like) loading surgical conditions for a late stage (P21) BT shunt procedure (black). Note the different scales from top to bottom rows. All values are reported as mean ± standard error of mean (*n* = 4-5 per group; see Table S1).

To simulate (upper bound) hemodynamic conditions for an ideal early BT shunt circuit (at P10, assuming perfect preservation of umbilical vessels), geometric and biomechanical metrics for E18.5 UV, E18.5 UA, and P10 RPA were also calculated at a (representative systolic) pressure of 15 mmHg and compared to P10 RSA at 56 mmHg (Tables 1 and S1). Axial stretch was maintained at *in vivo* values for RPA and RSA but set to unity for UV and UA, assuming that UV and UA would be implanted surgically at an unstretched length. The simulated pressure-distended inner radius increased most in E18.5 UV and slightly in P10 RPA, while it remained essentially unchanged in E18.5 UA, noting that these radii were each greater than in P10 RSA (Fig 2C). A similar simulated behavior was observed for circumferential wall stress, with elevated values in P10 RPA and E18.5 UV but decreased values in E18.5 UA at 15 mmHg (Fig 2D). All of these comparisons and calculations were repeated for an upper bound (representative systolic) pressure of 90 mmHg for P21 RSA and 20 mmHg for P21 RPA, E18.5 UA, and E18.5 UV to mimic an ideal shunt creation at P21 (i.e., with preserved umbilical properties), which revealed greater differences between the umbilical and postnatal vessels in both structural (Fig 2E) and material (Fig 2F) behaviors. Simulations for this later stage (P21) BT shunt procedure were nevertheless similar qualitatively (Fig 2G, H), with inner radius increasing significantly in UV and circumferential wall stress increasing most in P21 RPA and E18.5 UV compared to native conditions. UA experienced a mild increase in inner radius (Fig 2G) and non-significant reduction in circumferential stress (Fig 2H). Again, the simulations suggested that UV would experience greater changes than would UA in a simulated shunt condition. Overall, the umbilical vessels are better matched with RPA at P10 and RSA at P21 for the inner radius.

Findings for the native axial structural (Fig S1A, E) and material (Fig S1B, F) behaviors for *in vivo* conditions as well as for simulated ideal BT shunt procedures at P10 and P21 (Fig S1C, D, G, H) showed that axial stress reduced significantly in the umbilical vessels post-shunt due to implantation at an axial stretch of 1. Calculated values of axial stress and elastic energy stored upon biaxial deformation increased for RPA due to the simulated early (P10) and later (P21) BT shunt procedures, albeit not significantly. Together, these passive mechanical results suggest that UA may better tolerate the changing hemodynamics due to a BT shunt procedure, whereas both of the vessels normally subjected to lower pressures (PA and UV) would experience the most dramatic changes in wall stress and elastic energy storage due to deformation.

Noting that the period from P10 to P21 represents one of rapid growth of the postnatal murine vasculature (Murtada et al., 2021), maturation of native RPA from P10 to P21 appeared to associate with greater changes in most circumferential mechanical metrics relative to changes in RSA (Fig 2A, E; Table S1), with the latter showing consistently stiffer circumferential stress-stretch material responses (Fig 2B, F; Table S1). The *in vivo* axial stretch also increased from P10 to P21 in both RSA (∼12%, from a mean of 1.47 to 1.64) and RPA (∼22%, from a mean of 1.23 to 1.53), as seen in Table S1. The *in vivo* axial stretch was also high in UV (∼1.55), which in part gave rise to the increased inner radius when analyzed at an unloaded length (simulated shunt procedure, Fig 2C, G). Reflective of the different *in vivo* functions, RSA had a greater elastic energy storage capacity (mean values of 20 and 44 kPa at P10 and P21) than did RPA (3 and 13 kPa at P10 and P21), with the storage capacity of UV and UA at E18.5 very low (mean values of 1 and 3 kPa, respectively), which increased marginally in the simulated shunt procedures for UV and decreased for UA.

UV and UA exhibited different vasoconstrictive responses when stimulated with 100 mM KCl at isobaric *in vivo* relevant pressures of 5 versus 25 mmHg (Table S3), verified through side-view (video capture) and cross-sectional (OCT) images of cannulated vessels in contracted states (Fig 3A, E). Histological images revealed the high cellularity of the vascular walls, particularly in the outer layer of UA. Both UV and UA exhibited strong vasoconstriction at 5 mmHg, higher than P10 RSA and P10 RPA (Fig 3I, J), with the lumen closing in UA while remaining patent in UV at the steady-state contracted state (Fig 3A, B, E, F). When contracted at 25 mmHg, UA maintained a high vasoconstrictive response similar to that of P21 RPA while UV exhibited a weak vasoconstriction closer to the magnitude observed in P21 RSA (Fig 3K, L). The contractile response in UA was higher at 25 than 5 mmHg, perhaps reflecting a peculiar structure-function relationship.

**Fig 3.**
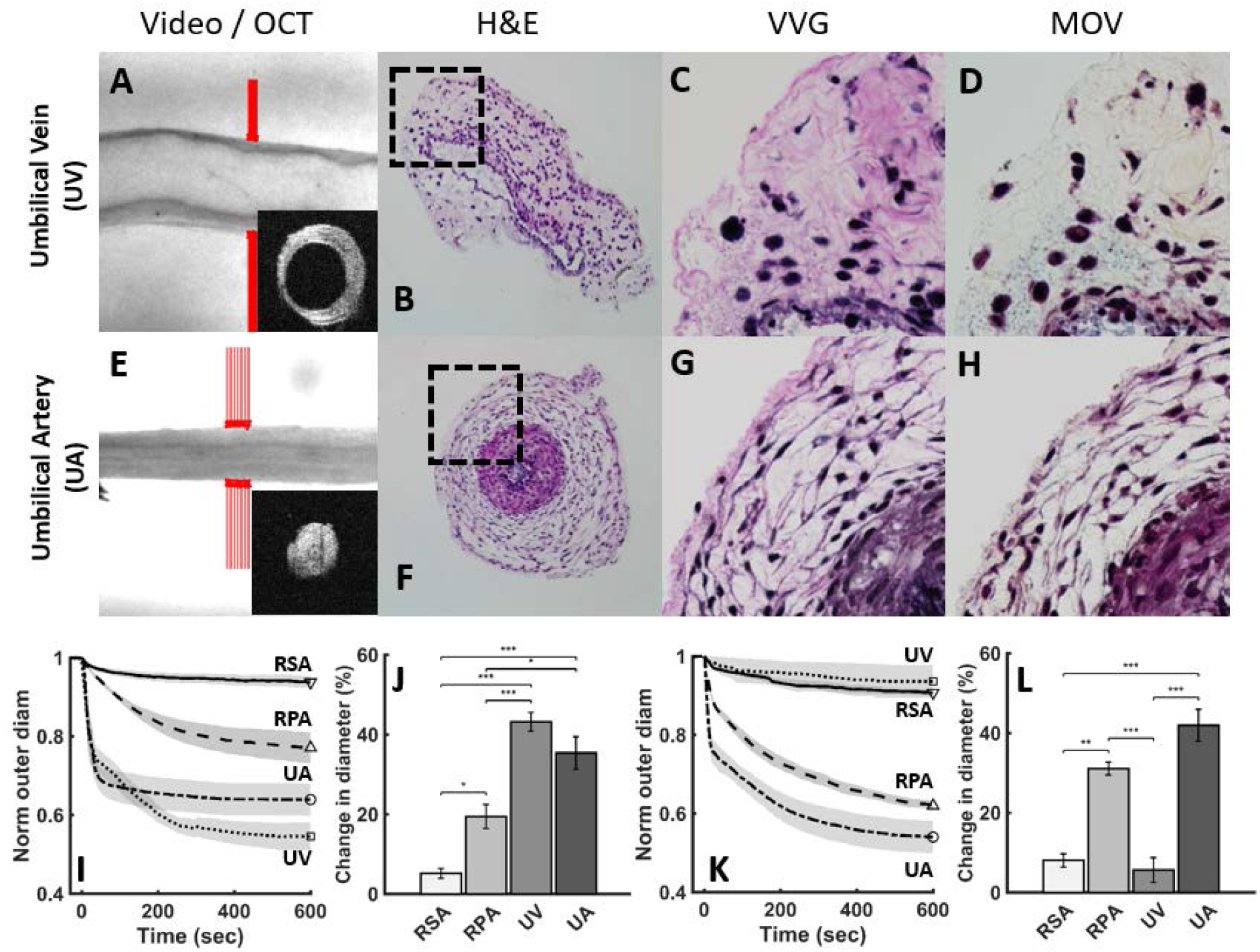
Regional vasocontractility of umbilical vein (UV) and umbilical artery (UA) at E18.5 and the right subclavian artery (RSA) and right pulmonary artery (RPA) at both P10 and P21. **A, E:** Side-view (video microscopy) and transverse cross-sectional (OCT, insets) images of UV and UA that were pressurized at 25 mmHg when stimulated with 100 mM KCl. See Fig S2 for similar results when stimulated with high KCl at 5 mmHg. **B-D, F-H:** Hematoxylin and eosin (H&E), Verhoeff–Van Gieson (VVG) and Movat’s Pentachrome (MOV) stain for UV and UA. **I, J:** Contractile responses at 5 mmHg were stronger in E18.5 UV (□) and UA (○) than in P10 RSA (▽, pressure = 30 mmHg) and RPA (Δ; pressure = 5mmHg). **K, L:** The UV was unable to vasoconstrict at 25 mmHg whereas the UA constricted well and closed its lumen at this pressure; there was also a weak vasoconstriction by the P21 RSA at 50 mmHg, whereas the P21 RPA constricted well at 7.5 mmHg. All values are reported as mean ± standard error of mean (*n* = 4-6 per group; Table S3).

Repeated contractions (>2 sequential KCl-induced contractions with intermediate Krebs washout-induced relaxations) of UV and UA resulted in bending after each relaxation (Fig 4A, D) as well as difficulties in distending the lumen fully (Fig 4B, E). Despite weak vasoconstrictive responses by UV at 25 mmHg, repeated KCl stimulations caused UV to experience constriction-related intramural damage (Fig 4C). Following repeated contractions, UA displayed dilatational difficulties as well, though with signs of inward buckling but less mural damage (Fig 4F).

**Fig 4.**
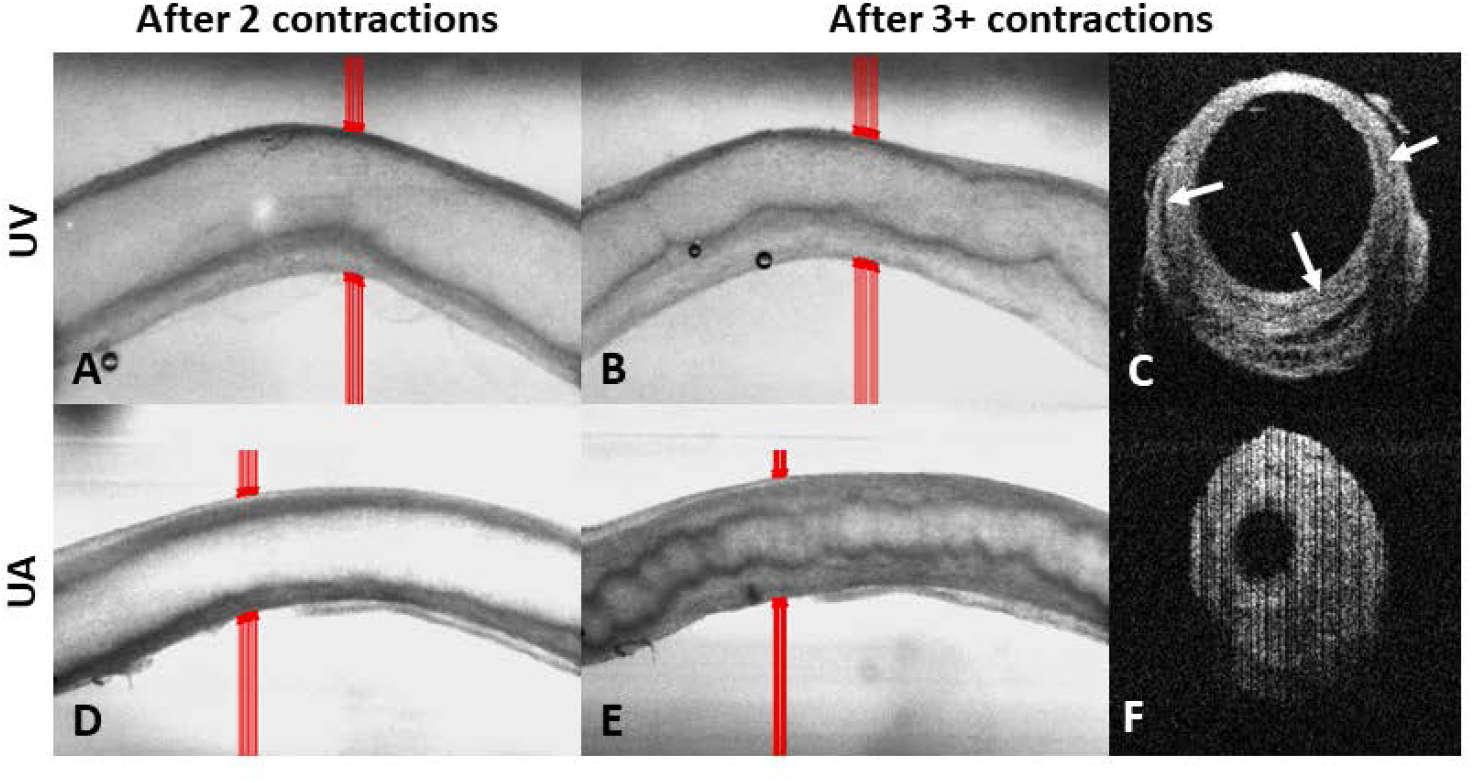
Sustained or repeated vasoconstrictions caused intramural mechanical damage in both the UV and UA. **A, B, D, E:** Longitudinal (video microscopy) and **C, F** transverse cross-sectional (OCT) images after two or more contractions reveal bending and reduced lumens in both umbilical vessels. Signs of more severe dissection-like damage was observed in the UV (*n* = 4-6 per group; Table S3). UA-umbilical artery, UV-umbilical vein, OCT -optical coherence tomography.

Finally, given concerns with repeated smooth muscle contractions and occluded lumen in UA at hypercontractile states, we decellularized additional UV and UA and performed passive biaxial testing. As expected, we observed a complete elimination of contractile response and a preserved intact vascular wall in the decellularized vessels. Decellularization decreased the luminal radius in both UA and UV (Fig 5A); it also resulted in a significant left-ward shift in the luminal pressure-radius (Fig S3A, B) and circumferential stress-stretch (Fig S3C, D) relationships but only a slight increase in (structural) compliance, not significant in either decellularized UA or UV (Fig 5B).

**Fig 5.**
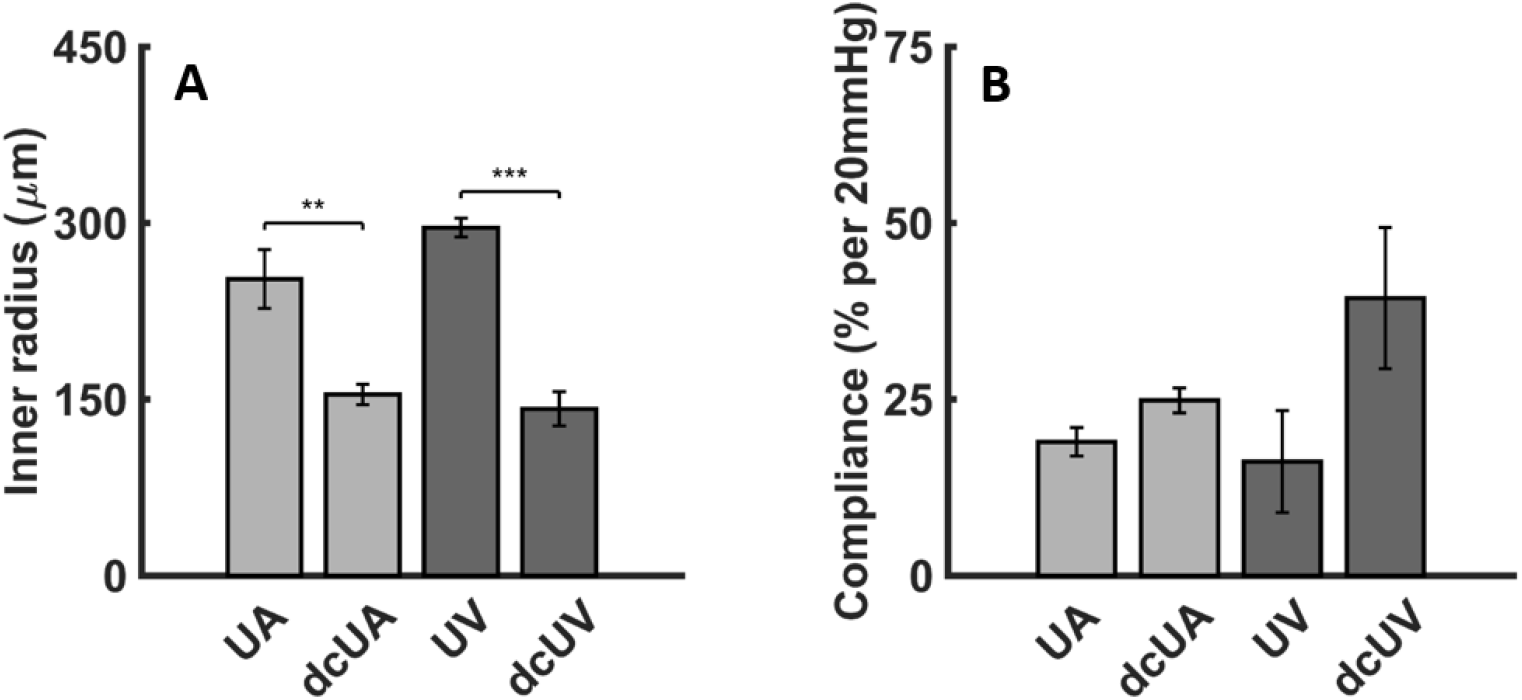
Geometric (luminal radius) and structural (compliance) metrics for the umbilical artery (UA) and umbilical vein (UV) either without or following decellularization (dc). Note that decellularization decreased the luminal radius in both vessels and tended to increase compliance in both, more so in the UV. All values are reported as mean ± standard error of mean (*n*=5 for dcUA, *n*=4 for all the other groups).

## Discussion

Despite frequent monitoring and use of anticoagulants to reduce the risk of thrombosis, current synthetic grafts used as BT shunts are not without complications (Petrucci et al., 2011), thus emphasizing a need to consider alternate surgical designs (Moghadam et al., 2012) and shunt materials (Hoganson et al., 2018). Selection of shunt materials (synthetic or native) should take into consideration the biaxial biomechanics, especially because all vascular cells are sensitive to changes in their local mechanical environment and may drive vascular remodeling if perturbed (Humphrey, 2008). The neonatal vasculature also undergoes significant changes in size and structure over a short period (Caspi et al., 2008; Ovroutski et al., 2009), thus constraining the use of non-growing synthetic shunts. Autologous umbilical vessels may thus serve as an alternative BT shunt material; they can also be used as access points for an external oxygenator for premature infants (Partridge et al., 2017). Yet, biomechanical characterization of UA and UV has remained limited (Dodson et al., 2013; Ferguson and Dodson, 2009).

Although the two umbilical vessels had similar inner radii at explant (250 μm for UV and 291 μm for UA), UA initially had a better matching simulated luminal radius (∼296 μm) relative to simulated *in vivo* P10 values for RSA (∼218 μm) and RPA (∼269 μm) than did UV (332 μm). Based on these simulated early BT shunt conditions, UA had a reduced circumferential wall stress (60% decrease) whereas UV had a markedly elevated (3.3-fold increase) circumferential wall stress. This stress tends to be highly mechano-regulated (Humphrey, 2008), hence decreases can lead to atrophy whereas increases can induce wall thickening, both over days. The simulated UV- and UA-BT shunts at P10 had marked decreases in axial stress (∼80% reduction for UV and ∼90% reduction for UA). A significantly reduced axial stress could also be detrimental to subsequent remodeling (Jackson et al., 2005), but this could be offset by an imposed axial stretch greater than unity at implantation or it could possibly correct itself due to somatic growth. Based on these and other simulated values of key geometric and passive biomechanical metrics, UV and UA could both offer potential advantages despite not being ideal.

Vessel-level contractility can play positive roles in modulating vascular remodeling (Dajnowiec and Langille, 2007; Valentin et al., 2009) and protective roles in otherwise structurally vulnerable vessels by carrying part of the hemodynamic load (Ferruzzi et al., 2016). However, the vasoactive nature of umbilical vessels has been reported to contribute to graft failure in newborn lambs (Vlessis et al., 1995). Here, we showed that contractile capacity was significantly higher in both UV and UA than in RSA and RPA at lower pressures, which could confer some advantages but also some concerns. First, the ability of UA to vasoconstrict against both a low (5 mmHg) and higher (25 mmHg) pressure, with complete luminal closure, suggests a need for care with regard to its use as a possible shunt material. By contrast, UV exhibited a much lower vasoconstrictive potential at the higher pressure (25 mmHg) and did not experience luminal closure even at the lower pressure (5 mmHg). Maintaining patency is critical to the success of a BT shunt, suggesting a possible advantage of UV over UA. Note that vasoconstrictive capacity was tested at fixed pressures different from the expected upper bounds at early (>15 mmHg) and later (>20 mmHg) BT shunt creation (Wang et al., 2020) to ensure consistency across tests. We did not pursue different pressures given that sustained and repeated vasoconstriction led to possible damage of the umbilical vessels, especially UV. The UA vasoconstricted spontaneously upon excision and both vessels experienced possible damage from repeated vasoconstriction, implying that it will be important to minimize contraction of umbilical vessels at the time of harvest.

Although a potential advantage of umbilical vessels as shunts is that they can be autologous, the potential concern with smooth muscle cell contraction and the need to preserve the vessels from the time of birth to the time of the first surgery led us to consider decellularization, which is increasingly common in tissue engineering (Crapo et al., 2011; Han et al., 2021). Following mechanical testing of the decellularized umbilical vessels, formalin-fixed paraffin-embedded H&E-stained sections were assessed to confirm the decellularization process but this was unsuccessful due to their small size. Nevertheless, biomechanical tests revealed a change in the circumferential stress-stretch response similar to that previously reported for successfully decellularized human umbilical arteries (Gui et al., 2009). Our pilot studies suggested further that compliance does not change dramatically with decellularization, although there was an unexpected reduction in luminal radius that could be a concern (Spigel et al., 2022). Further studies are warranted to compare different decellularization protocols.

Our computations also suggested that changes in wall mechanics could arise in RPA upon surgical creation of a BT shunt. Such changes could elicit adverse remodeling of the PA. Thus, one should consider methods of reducing BT shunt-induced pulmonary vascular remodeling, noting that reducing acute loading alone could have a benefit (Ramachandra et al., 2020). Such reductions could be achieved with external support of the pulmonary and similarly umbilical vessels used as shunts (Ramachandra et al., 2022). We did not experimentally provide or computationally simulate possible external support in any of the vessels, but this should be considered in the future. We also did not attempt to preserve the umbilical vessels from the time of harvest until the time of the simulated BT shunt procedure (10 or 21 days later). Rather we focused on ideal simulations wherein umbilical properties measured at the time of harvest (E18.5) were used to compute possible changes in mechanics if placed as shunts between RSA and RPA at either P10 or P21. The properties of the decellularized umbilical vessels were measured 5 days after harvest (i.e., after decellularization). There is a need for continued research into optimal methods of preservation of the umbilical vessels, but this was beyond the scope of the current mechanics-based simulations.

Finally, although the ultimate goal is to examine safety and efficacy of autologous umbilical vessels as BT shunts in humans, this study focused solely on murine studies. Human umbilical vessels are readily available for testing, but our murine studies enabled a critical new comparison – consistent quantification of biaxial biomechanical properties across four key vessels involved in a BT shunt procedure, including postnatal subclavian and pulmonary arteries that are not readily available from human donors. Notwithstanding differences between human and murine vessels, diverse mouse models continue to contribute significantly to our increasing understanding of vascular biology and mechanics. Indeed, murine data-informed computational models recently provided critical and consistent predictions of the in vivo development of tissue engineered vascular grafts in pre-clinical and clinical Fontan completion (Drews et al., 2020), hence providing further evidence of the utility of murine studies.

In summary, the potential use of umbilical vessels as autologous conduits has attracted the attention of both tissue engineers (Gui et al., 2009) and surgeons (Hoganson et al., 2018), but this report is the first to compare biaxial biomechanical properties across umbilical vessels and potential host vessels to which they will be anastomosed. If methods could be found to neutralize vessel-level contractility without compromising other actomyosin-mediated cell functions, including mechanosensing, UA and UV both appear to have potential for use as BT shunts, assuming that their properties can be preserved from the time of birth to the BT surgery. Otherwise, our data suggest that intact UV could be a better choice as a shunt than UA due to the risk of vasoactive lumen closure or mural damage. Decellularized UA may be a viable alternate option, however.

## Acknowledgment

This work was supported in part by Additional Ventures (AVCC). We would like to acknowledge Edward Han, PhD, for his assistance with the decellularization protocol and an anonymous reviewer who provided insightful suggestions that improved the clinical relevancy of this work.

