## Supplemental Materials for "*Ex Vivo* Biomechanical Characterization of Umbilical Vessels: Possible Shunts in Congenital Heart Palliation"

[2]Vascular Biology and Therapeutics Program
Yale School of Medicine, New Haven, CT, USA

[*]These authors contributed equally

**FIGURES**

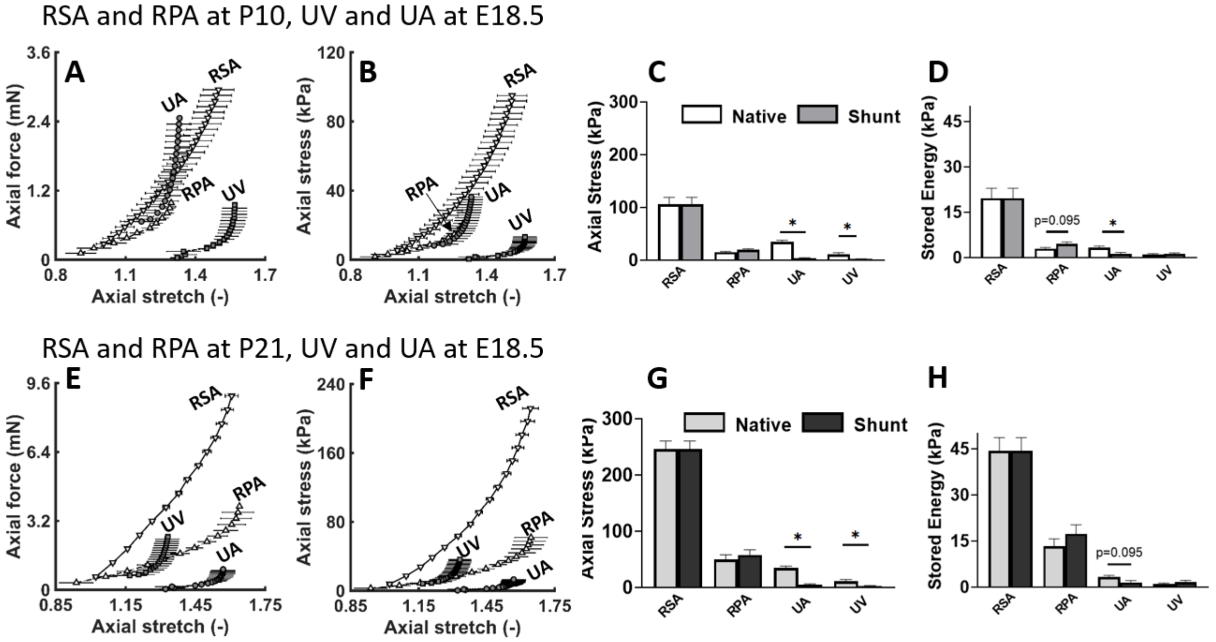


**Fig S1**. Comparison of regional passive biomechanical behaviors of the right subclavian artery (RSA, ▽) and right pulmonary artery (RPA, △) as well as the umbilical vein (UV, □) and umbilical artery (UA, ○). **A, B, E, F:** Similar to Figure 2 in the main text except for axial force-stretch and axial stress-stretch relationships. **A, B:** Relationships at postnatal day P10 for the RSA and RPA, with those for the UV and UA at embryonic day E18.5 shown for comparison. **E, F:** Relationships at P21 for the RSA and RPA, with those for the UV and UA at E18.5 again shown for comparison. **C, D**: Values of axial Cauchy stress and stored energy for vessel-specific physiological loading conditions for the RSA and RPA at P10 and the UV and UA at E18.5 (white bars) as well as at simulated (surgical-like) loading conditions for an early stage (P10) BT shunt procedure (dark grey). **G, H**: Similar to panels C, D except for the RSA and RPA at P21 and the UV and UA at E18.5 (light grey) as well as at simulated (surgical-like) loading conditions for a late stage (P21) BT shunt procedure (black). All values are reported as mean ± standard error of mean (*n* = 4-5 per group; see Table S1).

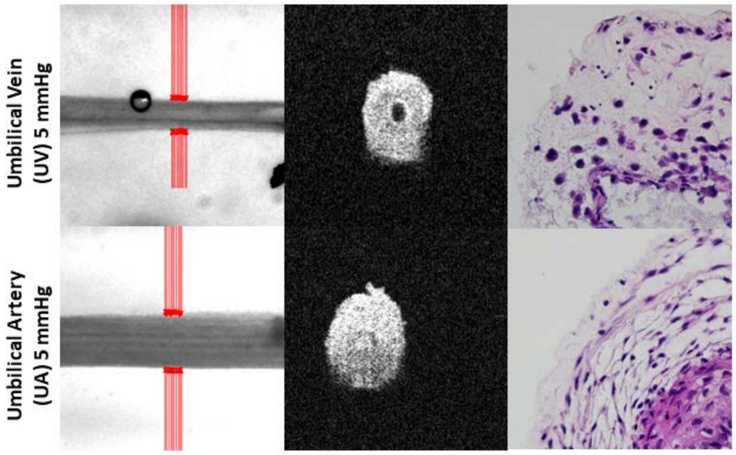


**Fig S2**. Similar to Figure 3 except for vasoconstriction of the umbilical vein (UV, top row) and umbilical artery (UA, bottom row) excised at E18.5, held at specimen-specific axial stretches, and pressurized to 5 mmHg before exposure to 100 mM KCl. Note that both vessels were able to reduce their lumen significantly, including closure of the lumen by the UA.

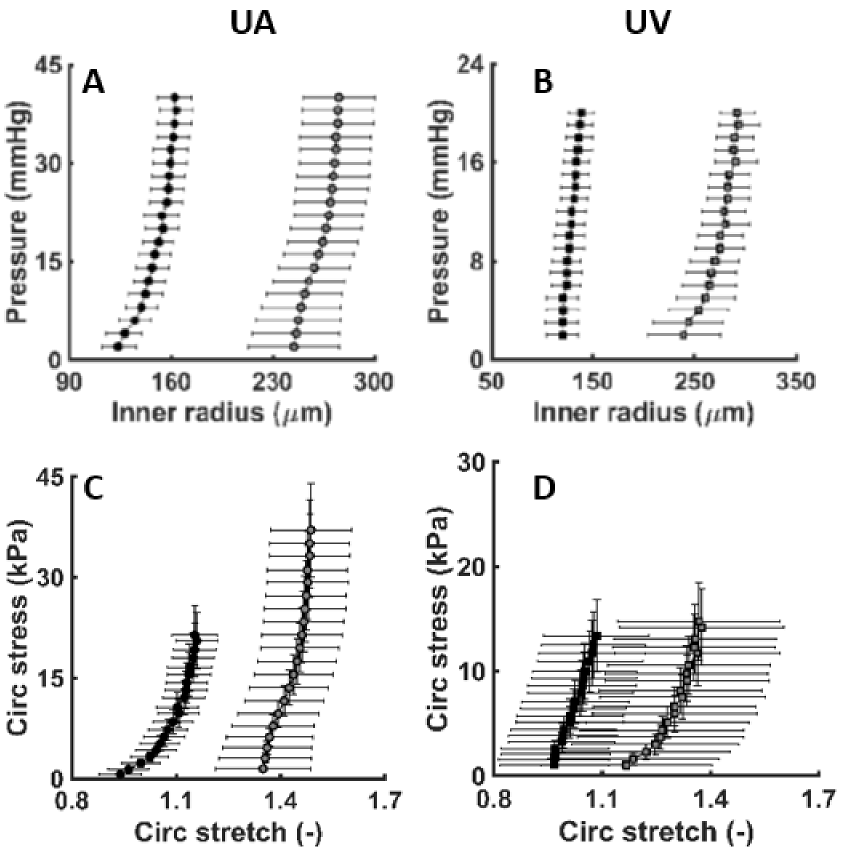


**Fig S3.** Pressure-inner radius (A, B) and circumferential stress-stretch relationships (C, D) of decellularized (solid) and intact (grey) umbilical arteries (left column) and umbilical veins (right column) near in vivo axial stretch. All values are shown as mean ± standard error of mean (*n*=5 for the decellularized umbilical artery, *n*=4 for all the other groups).

**Table S1**. Passive geometric and biomechanical metrics for the RSA and RPA separately at P10 and P21 and UV and UA at E18.5 derived from cyclic pressure-diameter and axial force-length testing while the smooth muscle was rendered relaxed. Shown, too, are values computed for a simulated BT shunt procedure performed at either P10 (pressure of 15 mmHg) or P21 (pressure of 20 mmHg). Many of these values were computed using parameter values found in Table S2. All values are reported as mean ± standard error of mean, with the number of samples (*n*) shown for each group.

| | **RSA-P10** | **RSA-P21** | **RPA-P10** | **After Shunt** | **RPA-P21** | **After Shunt** | **UA** | **After Shunt** | **After Shunt** | **UV** | **After Shunt** | **After Shunt** |
|---|---|---|---|---|---|---|---|---|---|---|---|---|
| | **n = 5** | **n = 5** | **n = 5** | | **n = 4** | | **n = 5** | | | **n = 4** | | |
| **Unloaded dimensions** | | | | | | | | | | | | |
| Wall Thickness (μm) | 54.38 ± 0.84 | 69.77 ± 1.02 | 59.04 ± 1.86 | | 61.02 ± 1.85 | | 75.96 ± 5.64 | | | 80.42 ± 12.09 | | |
| Outer Diameter (μm) | 348.06 ± 27.48 | 434.24 ± 11.33 | 392.22 ± 20.36 | | 638.59 ± 22.34 | | 522.00 ± 49.14 | | | 555.59 ± 72.43 | | |
| Axial Length (mm) | 1.49 ± 0.16 | 1.37 ± 0.13 | 1.30 ± 0.07 | | 1.95 ± 0.25 | | 3.60 ± 0.27 | | | 3.94 ± 1.05 | | |
| **Loaded dimensions** | **P = 56** | **P = 90** | **P = 11** | **P = 15** | **P = 17** | **P = 20** | **P = 30** | **P = 15** | **P = 20** | **P = 5** | **P = 15** | **P = 20** |
| Outer Diameter (μm) | 484.81 ± 13.07 | 540.14 ± 16.68 | 594.68 ± 19.40 | 642.05 ± 21.62 | 903.02 ± 33.40 | 984.37 ± 35.74 | 664.52 ± 65.48 | 701.45 ± 74.52 | 715.17 ± 77.02 | 586.86 ± 37.88 | 765.90 ± 25.99 | 783.78 ± 18.28 |
| Wall Thickness (μm) | 23.98 ± 2.70 | 30.56 ± 0.77 | 28.05 ± 1.40 | 25.81 ± 1.36 | 26.58 ± 1.88 | 24.27 ± 1.84 | 41.10 ± 4.70 | 54.74 ± 8.33 | 53.46 ± 8.08 | 43.92 ± 6.75 | 50.83 ± 7.19 | 49.49 ± 7.19 |
| Inner Radius (μm) | 218.43 ± 5.22 | 239.52 ± 8.95 | 269.29 ± 10.09 | 295.22 ± 11.27 | 424.93 ± 17.55 | 467.92 ± 18.87 | 291.16 ± 34.88 | 295.99 ± 41.68 | 304.12 ± 42.74 | 249.51 ± 22.17 | 332.12 ± 13.50 | 342.40 ± 9.68 |
| in vivo Axial Stretch | 1.47 ± 0.06 | 1.64 ± 0.05 | 1.23 ± 0.02 | 1.23 ± 0.02 | 1.53 ± 0.08 | 1.53 ± 0.08 | 1.35 ± 0.04 | 1.00 ± 0.00 | 1.00 ± 0.00 | 1.55 ± 0.05 | 1.00 ± 0.00 | 1.00 ± 0.00 |
| in vivo Circumferential Stretch | 1.63 ± 0.17 | 1.40 ± 0.05 | 1.72 ± 0.09 | 1.87 ± 0.10 | 1.52 ± 0.01 | 1.66 ± 0.01 | 1.41 ± 0.11 | 1.46 ± 0.12 | 1.49 ± 0.12 | 1.21 ± 0.14 | 1.62 ± 0.23 | 1.68 ± 0.26 |
| **Systolic Cauchy Stresses (kPa)** | | | | | | | | | | | | |
| Circumferential | 72.46 ± 9.91 | 94.49 ± 5.71 | 14.24 ± 1.01 | 23.19 ± 1.68 | 36.64 ± 3.46 | 52.51 ± 5.15 | 30.50 ± 5.33 | 12.38 ± 2.77 | 17.31 ± 3.83 | 4.19 ± 0.95 | 13.88 ± 2.05 | 19.59 ± 2.74 |
| Axial | 106.30 ± 12.74 | 245.84 ± 14.69 | 15.51 ± 1.31 | 19.75 ± 1.75 | 50.19 ± 8.21 | 58.16 ± 8.99 | 34.92 ± 2.93 | 3.61 ± 1.14 | 4.69 ± 1.61 | 10.86 ± 3.51 | 2.19 ± 0.44 | 3.07 ± 0.45 |
| **Systolic Linearized Stiffness (MPa)** | | | | | | | | | | | | |
| Circumferential | 0.53 ± 0.06 | 0.56 ± 0.04 | 0.09 ± 0.01 | 0.15 ± 0.02 | 0.14 ± 0.01 | 0.21 ± 0.02 | 0.48 ± 0.12 | 0.18 ± 0.05 | 0.30 ± 0.09 | 0.07 ± 0.02 | 0.29 ± 0.11 | 0.43 ± 0.17 |
| Axial | 0.69 ± 0.07 | 1.93 ± 0.17 | 0.14 ± 0.01 | 0.17 ± 0.01 | 0.25 ± 0.04 | 0.30 ± 0.05 | 0.91 ± 0.17 | 0.03 ± 0.01 | 0.04 ± 0.02 | 0.44 ± 0.35 | 0.02 ± 0.00 | 0.02 ± 0.00 |
| **Systolic Stored Energy (kPa)** | 19.64 ± 3.29 | 44.37 ± 4.34 | 3.04 ± 0.33 | 4.58 ± 0.59 | 13.28 ± 2.40 | 17.36 ± 2.90 | 3.30 ± 0.53 | 1.26 ± 0.49 | 1.58 ± 0.54 | 0.98 ± 0.33 | 1.22 ± 0.39 | 1.66 ± 0.57 |

**Table S2**. Mean passive mechanical model parameters for the four-fiber family stored energy function. All values are reported as mean ± standard error of mean ($n$ = 4-5 per group; see Table S1).

| | | Elastic fibers | Axial Collagen | | Circ. Coll +SMC | | Symmetric diagonal collagen | | | Error |
|---|---|---|---|---|---|---|---|---|---|---|
| | | c (kPa) | $c_1^1$ (kPa) | $c_2^1$ | $c_1^2$ (kPa) | $c_2^2$ | $c_1^{3,4}$ (kPa) | $c_2^{3,4}$ | $\alpha_O$ (deg) | RMSE |
| RSA | P10 | 0.21 | 26.12 | 1.0E-09 | 7.31 | 1.0E-09 | 5.60 | 0.86 | 43.92 | 0.09 |
| | P21 | 12.16 | 29.28 | 0.01 | 17.62 | 1.0E-09 | 3.65 | 1.23 | 39.21 | 0.10 |
| RPA | P10 | 0.39 | 4.13 | 2.11 | 0.69 | 0.12 | 2.83 | 0.70 | 34.44 | 0.10 |
| | P21 | 5.73 | 1.32 | 4.65 | 2.41 | 1.0E-09 | 6.07 | 0.21 | 36.67 | 0.11 |
| UA | E18.5 | 1.99 | 0.25 | 12.84 | 1.59 | 1.66 | 1.30 | 5.10 | 40.01 | 0.25 |
| UV | E18.5 | 0.68 | 0.11 | 3.95 | 0.24 | 0.04 | 1.30 | 8.08 | 49.86 | 0.48 |

**Table S3**. Outer diameter and circumferential wall stress of P10 and P21 RSA and RPA and E18.5 UV and UA based on smooth muscle contraction with 100 mM KCl during biaxial isobaric – axially isometric tests. All values are reported as mean ± standard error of mean, with the number of samples (*n*) shown for each group.

| | RSA | | RPA | | UV | | UA | |
|---|---|---|---|---|---|---|---|---|
| | **P10** | **P21** | **P10** | **P21** | **5 mmHg** | **25 mmHg** | **5 mmHg** | **25 mmHg** |
| | **n = 5** | **n = 4** | **n = 5** | **n = 4** | **n = 4** | **n = 4** | **n = 6** | **n = 6** |
| **Loaded configuration** | | | | | | | | |
| Relaxed outer diameter (µm) | 408 ± 23.6 | 478 ± 6.2 | 470 ± 47.7 | 693 ± 25.2 | 583 ± 48.5 | 666 ± 22.3 | 568 ± 36.8 | 633 ± 34.3 |
| Contracted outer diameter (µm) | 388 ± 25.4 | 439 ± 8.4 | 374 ± 28.2 | 478 ± 21.4 | 330 ± 23.1 | 635 ± 40.8 | 363 ± 25.3 | 364 ± 25.6 |
| Change in outer diameter (%) | -5 ± 1.3 | -8 ± 1.7 | -19 ± 3.0 | -31 ± 1.6 | -43 ± 2.3 | -5 ± 3.5 | -35 ± 4.1 | -42 ± 4.0 |
| **Stress calculations** | | | | | | | | |
| Relaxed circumferential stress (kPa) | 19 ± 1.0 | 29 ± 1.4 | 3 ± 0.7 | 7 ± 0.4 | 2 ± 0.8 | 17 ± 2.6 | 2 ± 0.6 | 16 ± 3.3 |
| Contracted circumferential stress (kPa) | 17 ± 0.6 | 23 ± 1.1 | 2 ± 0.4 | 2 ± 0.3 | | | | |
| Change in circumferential stress (%) | -13 ± 3.1 | -21 ± 4.1 | -49 ± 5.5 | -65 ± 3.3 | | | | |
